# Pose estimation-based visual perception system for analyzing fish swimming

**DOI:** 10.1101/2022.09.07.507033

**Authors:** Xin Wu, Jipeng Huang, Lianming Wang

## Abstract

Advances in modern deep learning-based computer vision perception techniques have revolutionized animal movement research methods. These techniques have also opened up new avenues for studying fish swimming. To that end, we have developed a visual perception system based on pose estimation to analyze fish swimming. Our system can quantify fish motion by 3D fish pose estimation and dynamically visualize the motion data of marked keypoints. Our experimental results show that our system can accurately extract the motion characteristics of fish swimming, which analyze how fish bodies and fins work together during different swimming states. This research provides an innovative idea for studying fish swimming, which can be valuable in designing, developing, and optimizing modern underwater robots, especially multi-fin co-driven bionic robotic fish. The code and dataset are available at https://github.com/wux024/AdamPosePlug.

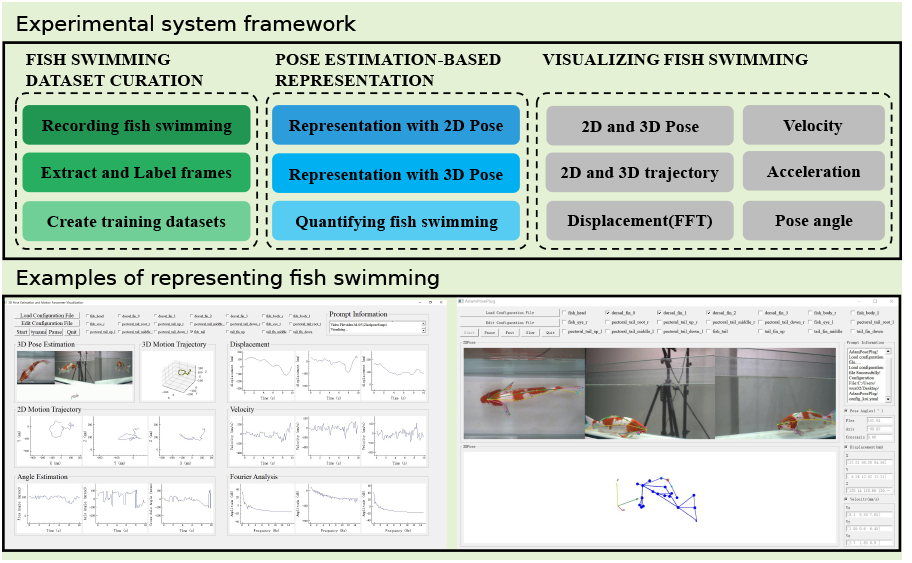

## I. Introduction

**A**dvanced deep learning-based computer vision perception techniques have facilitated the rapid development of animal behavior analysis tools that have greatly enhanced the ability of neuroscience and zoology scientists to conduct relevant research [1–3]. Among the behaviors of many species, we are more interested in the swimming of fishes because studying the swimming of fishes may reveal inspiring knowledge valuable for developing new types of robotic fishes [4].

Fish have evolved over millions of years to develop highly efficient locomotor skills that far exceed traditional engineered marine locomotion systems. Their excellent maneuverability and coordination inspire engineers to design and control novel underwater robots within a bionic framework [5–7]. The secret of efficient swimming in fish contributes to human understanding of fish behavior, physiology, and ecology and, at the same time, provides essential data and theoretical support for human design and control of novel underwater robots. Therefore, constructing a key that can unlock the secret of efficient fish swimming has become an important research direction. Recognition, detection, and measurement techniques based on advanced computer vision technologies have been developed recently. Researchers have constructed many fish motion analysis systems based on computer vision technology.

### 2D visual perception system for fish swimming

Jing et al. [8] recorded the motion of fish using video and studied the turning kinematics of a frightened carp. Li et al. [9] built on the previous research on the dorsal fin of carp and recorded richer 2D motion parameters. Yan et al. [10] designed observation systems that can track the two-dimensional motion of fish more accurately. However, these systems mainly acquired or followed the fish swimming condition in pixel space and needed to realize three-dimensional movement observation to extract more practical features.

### 3D visual perception system for fish swimming

Wu [11] constructed a video tracking system for 3D kinematics of fish using two cameras as well as a reflector, which was able to be used to extract ventral and lateral views of the fish, and with which the pitching and tailing motions of the fish were investigated. Zhang et al. [12] used X-ray photography to determine the number and length of vertebral joints, which supported the design of joint control for a carp bionic robot. Wang et al. [13] used a high-speed camera system to analyze the three-dimensional motion of a koi carp, focusing on its pectoral fins. Voesenek et al. [14] proposed a method for tracking the three-dimensional swimming of fish using multiple high-speed cameras designed with a video system based on image morphological transformations. Mao et al. [15] discussed the basic techniques of unobstructed tracking, investigated the method of unobstructed fish tracking, and realized the three-dimensional unobstructed fish tracking. Saberioon and Cisar [16] developed a three-dimensional occlusion-free search of fish using the structured light sensor. Qian and Chen [17] built a simple and cost-effective 3D motion detection device for zebra fish, enabling researchers to track the 3D motion of multiple zebra fish at low cost and precise spatial locations. Cheng et al. [18] proposed a method to acquire quantitative 3D motion trajectories of individual fish using a synchronized camera. These methods accept more information than 2D vision systems. However, they are limited by traditional computer vision methods, resulting in the extraction and understanding of localized, complex motion features that still need to be improved. Liu et al. [19] earlier proposed a technique for three-dimensional fish tracking using skeleton analysis and acquired the three-dimensional trajectories of fish. However, it was still limited to traditional vision methods and did not acquire more features.

These fish swimming observation systems mainly focus on 2D/3D movement tracking and simple movement pattern analysis. However, due to the fish’s disorganized movements, these methods cannot comprehensively analyze localized complex motions, such as detailed characterization of the caudal or pectoral fins. Therefore, extracting local complex motion features of fish using traditional computer vision methods is challenging, and a more effective tool is needed to remove more information to analyze local complex motion.

Recently, the rapid development of deep learning-based computer vision technology has made it an efficient way of solving complex feature extraction problems [20–22]. Deep learning models can quickly obtain the position and pose of the target object in the target coordinate system. Numerous advanced visual inference tools for animal behavior analysis [23–27] have been developed, and we also report on an algorithm for characterizing the three-dimensional motion of a target [28]. These advanced computer vision algorithms provide an essential theoretical basis for developing a visual system for observing fish swimming.

In this study, we constructed an visual perception system for studying fish swimming based on pose estimation. This advanced computer vision technique provides a new method for the study of fish swimming and is expected to be further applied to the analysis of complex movements of fish. The main research contents and contributions include the following three aspects:

1. We design an visual perception system for fish swimming research centered on a multi-view visual perception system for collecting fish swimming, equipped with a high-speed camera, a high-performance computer, complementary light, and flow generation. Thanks to the implementation of multi-view visual perception and advanced animal pose estimation algorithms, this system represents a significant improvement over others. It can extract a broader range of motion features, such as 2D/3D pose estimation, 2D/3D motion trajectories at crucial points, displacement, velocity, acceleration, and frequency. As a result, researchers can gain a more comprehensive understanding of the intricate dynamics involved in fish swimming.
2. We design the fish swimming visualization software (AdamPosePlug) based on animal pose estimation. The software integrates four functions of fish swimming data preprocessing, pose estimation, motion feature extraction, and visualization for easy use by researchers.
3. We use the visual perception system to study a variety of motions (advancing, retreating, turning, and floating up.) of koi in the tank. The experimental results show that our system can accurately extract the motion features of fish swimming. Moreover, with the help of these motion features, the synergy between the fish body and fins can be analyzed in detail, which may have some reference significance for the development of multi-fin co-driven bionic robotic fish.

The remainder of this paper is presented as follows: Section 2 describes the design of the visual perception system for studying fish locomotion. Section 3 presents the results of applying the visual perception system to study the koi movement and discusses the strengths of the practical system and details to be improved. Finally, Section 4 provides the concluding remarks of this research.

## II. Materials AND METHODS

In this section, we describe the technical details of our proposed framework.

### A. Overall framework

To record videos of fish swimming comprehensively and represent fish swimming in more detail, we constructed an visual perception system for studying fish swimming based on pose estimation. The practical system integrates a video capture device designed to quickly record fish swimming, an advanced algorithm to represent fish swimming and an easy-to-use visualization software.

The overall experimental framework is depicted in Figure 1. This consists of four main stages: (1) Fish swimming dataset curation (Section II-B); (2) Pose estimation-based representation (Section II-C); (3) Quantifying fish swimming (Section II-D); and (4) Visualizing fish swimming (Section II-D).

**Fig. 1.**
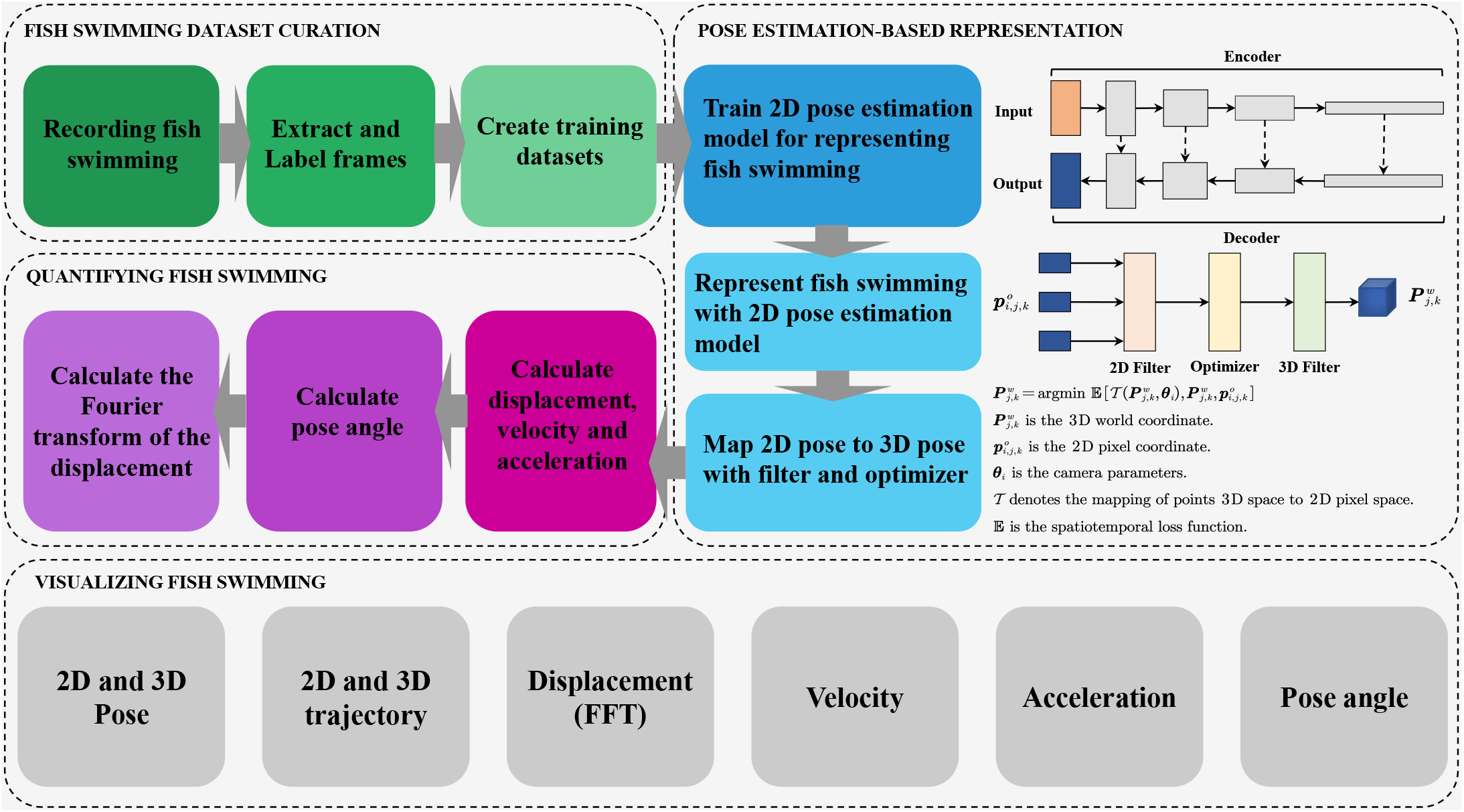
The proposed experimental framework consists of four phases. The dashed box includes four different phases, each of which, in turn, contains different steps. The first phase is fish swimming dataset curation, which mainly includes data acquisition, validation, proper extraction and labeling, and creating a dataset suitable for use in the second phase. The second phase is pose estimation-based representation, which mainly includes 2D pose estimation model training, using the 2D pose estimation model, and mapping 2D poses to 3D poses. The third phase quantifies fish swimming, mainly involving calculating various motion parameters. The fourth stage is visualizing fish swimming, which includes six visualization modules.

### B. Fish swimming dataset curation

The platform consists of the following equipment: a water tank (120 cm × 20 cm × 30 cm in length, width, and depth, respectively), a high-performance computer, three high-frame-rate (the maximum of frame rate is 210 FPS), and high-resolution (1280 × 1024 pixels) CCD cameras, a wave maker, and two fill lights (the maximum of power is 100 W). The schematic diagram of the data acquisition equipment is shown in Figure 2(a).

**Fig. 2.**
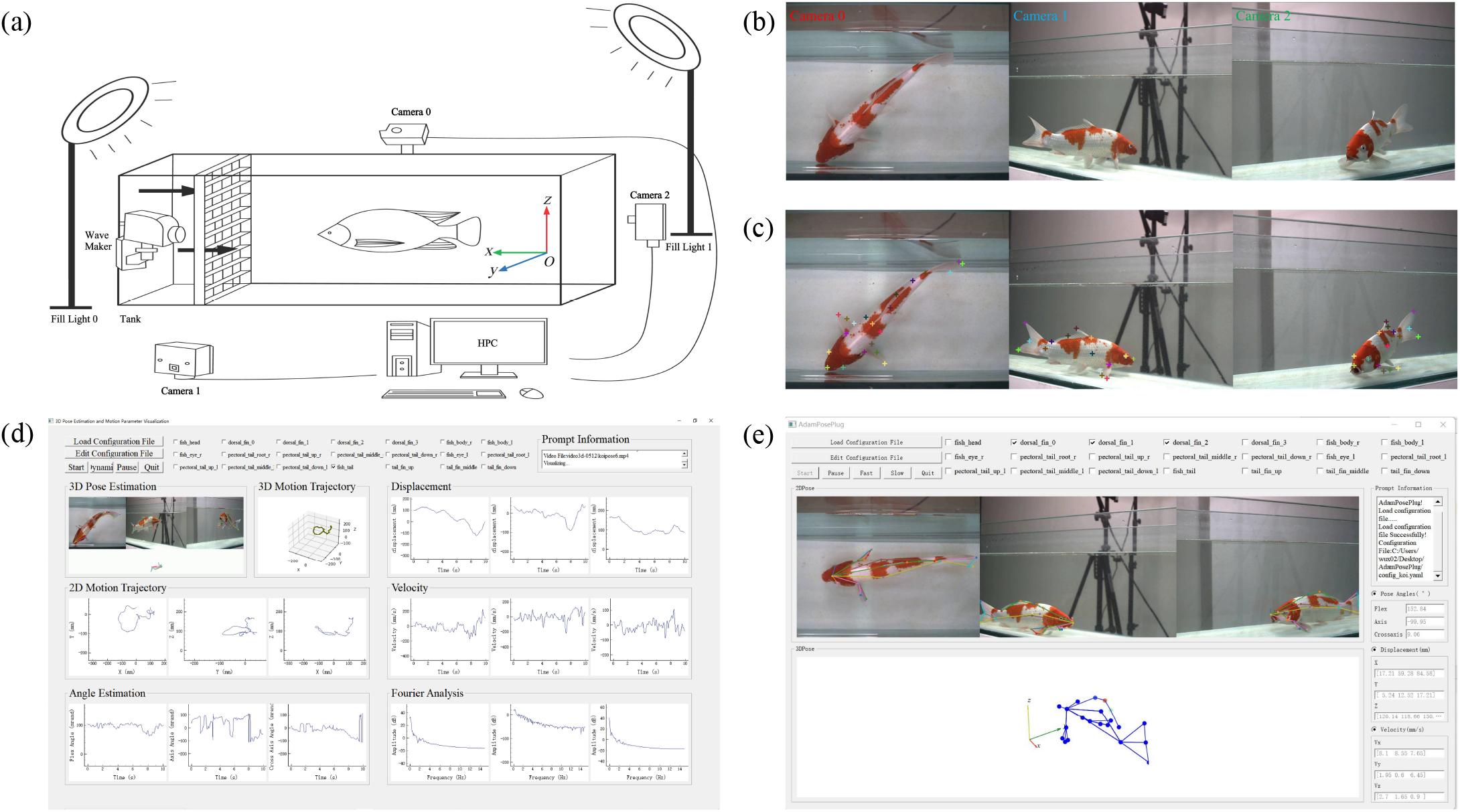
(a) This is a schematic of the fish swimming research system; (b) an example of image acquisition using the visual perception system; (c) an example diagram of koi key point labeling; (d)(e) and the software for quantifying and visualizing fish swimming. We shared an example of the visualization software run on [Google Drive](https://drive.google.com/file/d/1NhB6-OZ-CEMPxHnpZFEsT2DmufwGIU4C/view).

We control the capture device to record the movement of the fish in the pool from three different viewpoints simultaneously (more views can be extended). In addition, the wavemaker could simulate some typical water currents, and the fill light could provide even sunlight for the recording process. Figure 2(b) shows the fish swimming images recorded simultaneously from different views by the capture device.

In addition, we need to produce fish swimming datasets that can be used to train 2D pose estimation models. We marked the koi carp with customized 21 keypoints, including fish head (1), fish eyes (1 × 2), dorsal fin (4), pectoral fin (4 × 2), fish body (1 × 2), and caudal fin (4). Figure 2(c) shows an example of labeled koi. We also shared all of our data on Google Drive for researchers.

### C. Pose estimation-based representation

The remarkable performance when using deep learning for human 2D and 3D pose estimation plus dense representations made this large body of work ripe for exploring its utility in animal behavior. Many machine learning-based tools for laboratory experiments have arrived in the past five years. However, these methods are imperfect for 3D pose representation, so we designed algorithms [28] to map 2D pose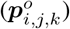 to 3D pose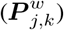.

#### 1) 2D pose representation for fish swimming

As quality tools for 2D animal pose estimation, DeepLabCut [23] and SLEAP [26] show different strengths, respectively; DeepLabCut supports more baseline models and is more accurate, while SLEAP is lighter and can adapt to high frame rates while maintaining accuracy. In addition, they still have in common that both use the encoder-decoder model structure.

DeepLabCut [23, 25] supports backbone including ResNet [29], MobileNet [30–32], EfficientNet [33], and DLCNet [25]. SLEAP [26] supports backbone networks including LEAPCNN [24], U-Net, Hourglass [34], and ResNet [29].

We used ResNet-152 as a baseline model to train models by DeepLabCut that can represent the 2D pose of a fish. At the same time, the encoder and decoder use skip connection [29] to avoid gradient vanishing or explosion during training. The encoder plays the role of feature extraction, while the decoder plays the role of feature re-representation. The encoder mainly consists of convolution (downsampling) kernels, while the decoder consists of deconvolution (upsampling) kernels of the corresponding dimensions. We use SGD [35] as an optimizer during training.

#### 2) 3D pose representation for fish swimming

After we obtain the 2D pose representation of the fish, we can map the 2D pose to the 3D pose by solving the optimization problem (1).

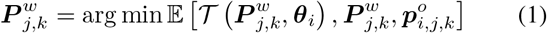

In the formula (1),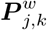 is the 3D world coordinate, 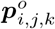 is the 2D piexl coordinate, ***θ***_*i*_ is the camera parameters, 𝒯 denotes the mapping of points 3D space to 2D piexl space, and 𝔼 is the spatiotemporal loss function. The algorithm we designed [28] knows all the specific details of solving the optimization problem.

In summary, we have implemented a 3D pose representation of fish swimming by combining DeepLabCut and the designed optimization algorithm. In the next stage, we can quantify the fish swimming by calculating the 3D pose representation.

### D. Quantifying and visualizing fish swimming

Computing on the time series composed of 3D keypoints, the displacement, velocity, and acceleration of each key point can be obtained, which are the most basic features to characterize fish swimming.

Tracking any one or more of the 2D keypoints and 3D keypoints can get the trajectory of the target in 2D-pixel space and 3D real space, which is the most essential feature representing the behavior of the fish.

Fourier analysis of the time series composed of displacements can analyze the possible periodic motion of the fish swimming.

In addition, the angles constituted by any three keypoints can be calculated based on the geometric relationship, and these angles can represent the local (fin) bending of the fish, which provides a new perspective for the analysis of the local complex motion of fish.

To facilitate easy and quick access to these motion features and to better present these results to the experimenter, we designed software (AdamPosePlug) to integrate these steps, and the software code has been open-sourced on GitHub.

The software runs as shown in Figure 2(d) and Figure 2(e). The interface contains the six modules in the fourth stage. The software is divided into two primary interfaces, which mainly qualitatively display the 2D and 3D postures of the fish swimming and quantitatively display the various motion characteristics of the fish swimming, thus facilitating the experimentalists to analyze the fish swimming. The main interface displays the motion parameters that show changes over time, as well as the 2D pose and 3D pose of the fish. The plug-in interface shows more prominently the 2D pose and 3D pose of the fish, as well as displaying motion parameters for selected keypoints.

## III. Results AND DISCUSSIONS

This section presents the results of our experiments using the visual perception system to characterize and visualize koi movement in the experimental tanks and provides some insights into koi movement. Figure 3 illustrates the actual visual perception system, and Table I lists the main parameters of the visual perception system.

**TABLE I.** Equipment parameter.

|  |  |
| --- | --- |
| Camera Model | OSG130-210UC |
| Color/Gray | Color |
| Pixel size | $4.8\mu\text{m} \times 4.8\mu\text{m}$ |
| Effective Pixels | $1280 \times 1024$ |
| Frame Rate Initial Value | 210 |
| Trigger Mode | Software Trigger |
| Data Interface | USB3.0 |
| Computer Model | Dell |
| Processor | Intel® Xeon SP® 6248R@3GHz |
| Machine with RAM | 128.0GB |
| GPU | RTX 3090 |
| System Type | X64-based Processor |
| Operating System | Windows 10 |

**Fig. 3.**
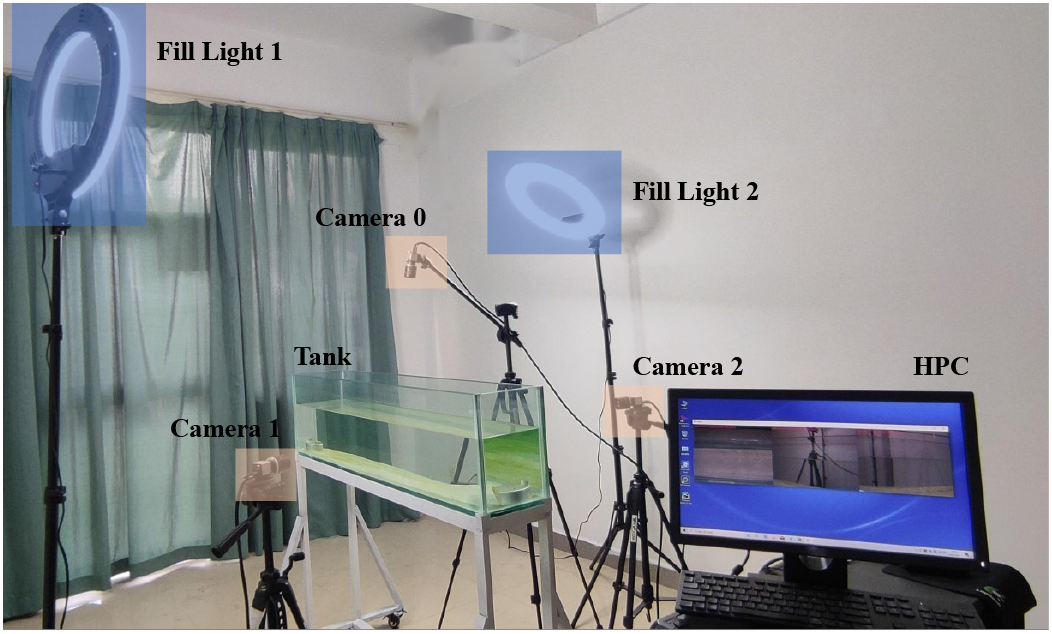
This is an visual perception system for studying fish swimming. The system differs somewhat from Figure 1(a). Since we have not conducted any research on fish swimming in complex flow fields for the time being, we did not install a wave maker. In addition, the placement of the high-speed camera is adjusted to collect information from all directions of fish swimming adequately. We also adjust the position and height of the fill light to give the visual perception system a more even and whole light.

The koi (300 mm) used in the experiment were kept in a tank. The experimental platform recorded 40 minutes of koi movement from three different angles, divided into 10-second segments, for a total of 240 videos. The videos were processed using software, and all 3D pose data obtained were stored on [Google Drive](https://drive.google.com/drive/folders/1c1kd9vjYOw07oB6Nz5_N-PGVi3h7-b8T?usp=drive_link). We extracted several segments of these data for analysis, and the experimental research was divided into five main parts:

1. We use a perceptual system to capture 2D and 3D keypoints of the fish to realize pose representation for fish swimming.
2. We use the keypoints extracted in step (1) to compute motion features such as motion trajectory, displacement, velocity and period, providing solid data support for subsequent fish swimming analysis.
3. We extracted five keypoints labeled with the dorsal fin at the top view angle for analyzing the fish’s advancing, retreating, and turning and showed 2D and 3D pose representations of its floating up.
4. We extracted 13 keypoints from the body, caudal, and pectoral fins to analyze koi’s body-fin coordination in four different swimming states (advancing, retreating, turn, and upward).
5. We extracted pectoral fin keypoints and then computed pose angles for analyzing pectoral fin flexion in koi under four different swimming states (advancing, retreating, turning, and upward) to understand the localized complex motion of the pectoral fins.

### A. Pose representation for fish swimming

To accurately analyze the movements of fish swimming, it is essential to utilize a perception system to obtain both 2D and 3D keypoints. Figure 4 serves as an example of how this can be achieved. We has developed a highly accurate 3D animal pose estimation algorithm [28].

**Fig. 4.**
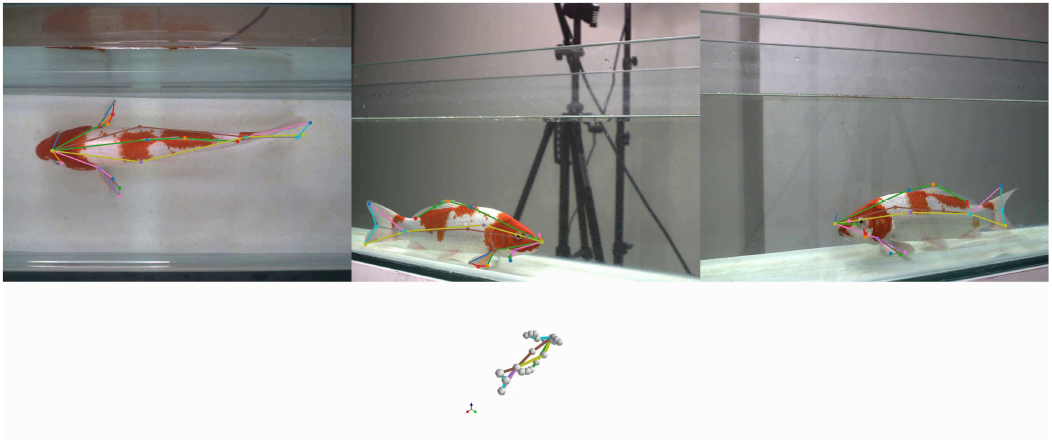
Pose representation for fish swimming. The first row shows the 2D pose representation for different viewpoints, and the second row shows the 3D pose representation.

99.83% of the keypoints reprojection error is less than 18 pixels, 93.41% of the keypoints reprojection error is less than 12 pixels, and 86.36% of the keypoints reprojection error is less than 5 pixels, demonstrating the precision of our approach. With this level of accuracy, we can better calculate the motion characteristics of the fish and gain deeper insights into their behavior.

### B. Quantifying fish swimming

Past approaches to analyzing fish swimming [11–19] have primarily relied on 2D/3D motion tracking and fundamental motion pattern analysis. However, these methods have yet to prove sufficient in capturing a comprehensive range of motion characteristics. To conduct a more thorough study of fish swimming, extracting additional motion parameters for analysis, such as motion trajectory, speed, and swing period is crucial. To illustrate our method, we demonstrate the extraction of motion parameters from koi swimming in a tank.

Take the keypoint at the upper end of the caudal fin as an example. The motion trajectory of this keypoint is shown in Figure 5. We have plotted the 3D motion trajectory of this keypoint and the 2D motion trajectory of this keypoint in the three planes of *xOy, yOz*, and *xOz*, and the direction of the target’s motion can be determined based on the 2D or 3D motion trajectory of the keypoint. In addition, it is possible to determine whether there is a periodic motion. When the fish is swimming, the swinging of the tail fin can be approximated as a periodic motion, and the clumped motion trajectories can determine that the visualization results are consistent with reality.

**Fig. 5.**
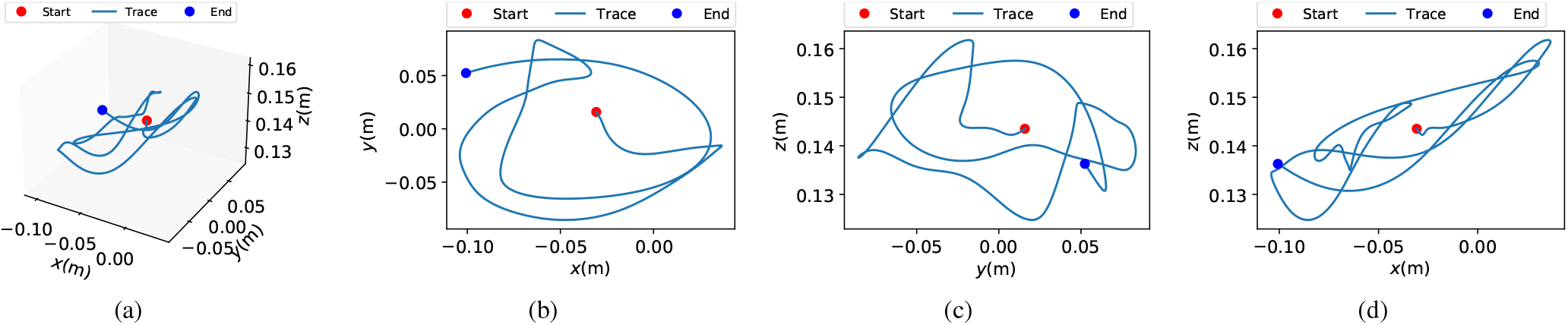
Motion trajectories of keypoints at the upper end of the tail. Movement trajectories help us track the behavior of fish swimming, especially periodicity. (a) 3D motion trajectory; (b) ***xOy*** plane 2D motion trajectory; (c) ***yOz*** plane 2D motion trajectory; and (d) ***xOz*** plane 2D motion trajectory. The red dots indicate the start of the keypoints, the blue dots indicate the end of the keypoints, and the blue curve indicates the motion trajectory.

We also obtained other motion features such as displacement, velocity, and the Fourier transform, where the Fourier transform can be used as a feature to analyze periodic or quasi-periodic motion. As seen from Figure 6(a), the motion includes a quasi-periodic motion with a period of about 4 ∼ 5s. Adult koi do not swing their caudal fins rapidly when undisturbed, so the period is more extended, but the swing amplitude is more significant. Figure 6(b) shows the change in velocity of keypoint motion over time. The velocity still contains a periodic variation. Without perturbation, the fish does not swim very fast, with a combined velocity of about 0 ∼ 150 mm/s. Figure 6(c) is the Fourier transform of the displacement. From the Fourier transform result, it can be seen that the second peak is 0.2322Hz at the peak point except for 0Hz, which means that the swing frequency at the key point of the caudal fin is about 0.2322Hz.

**Fig. 6.**
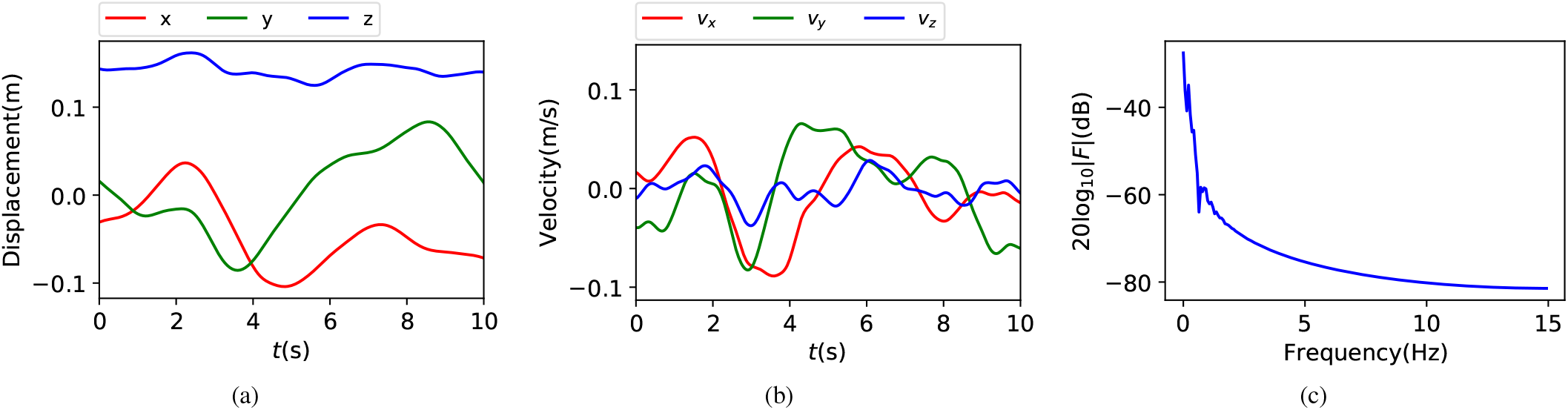
Motion parameters of keypoints at the upper end of the tail. Displacement, velocity, and Fourier analysis help us understand the various periodic movement patterns of the fish, which can provide data support for the design of bionic robotic fish. (a) Displacement; (b) Velocity; and (c) Fourier transform. Different colored curves in each graph indicate the displacement and velocity characteristics at different directions.

### C. Overlooking the koi swimming

This section focuses on extracting information about the keypoints of the dorsal fins of koi in the three swimming states of advancing, retreating, and turn in the top view angle. A total of 5 points, marked on the back of the fish at the locations of *x* = 0.11, 0.15, 0.17, 0.21, 0.30 m, are selected for the analysis. The *y*-direction displacements of these points are shown in Figure 7(a), plotted along the time axis.

**Fig. 7.**
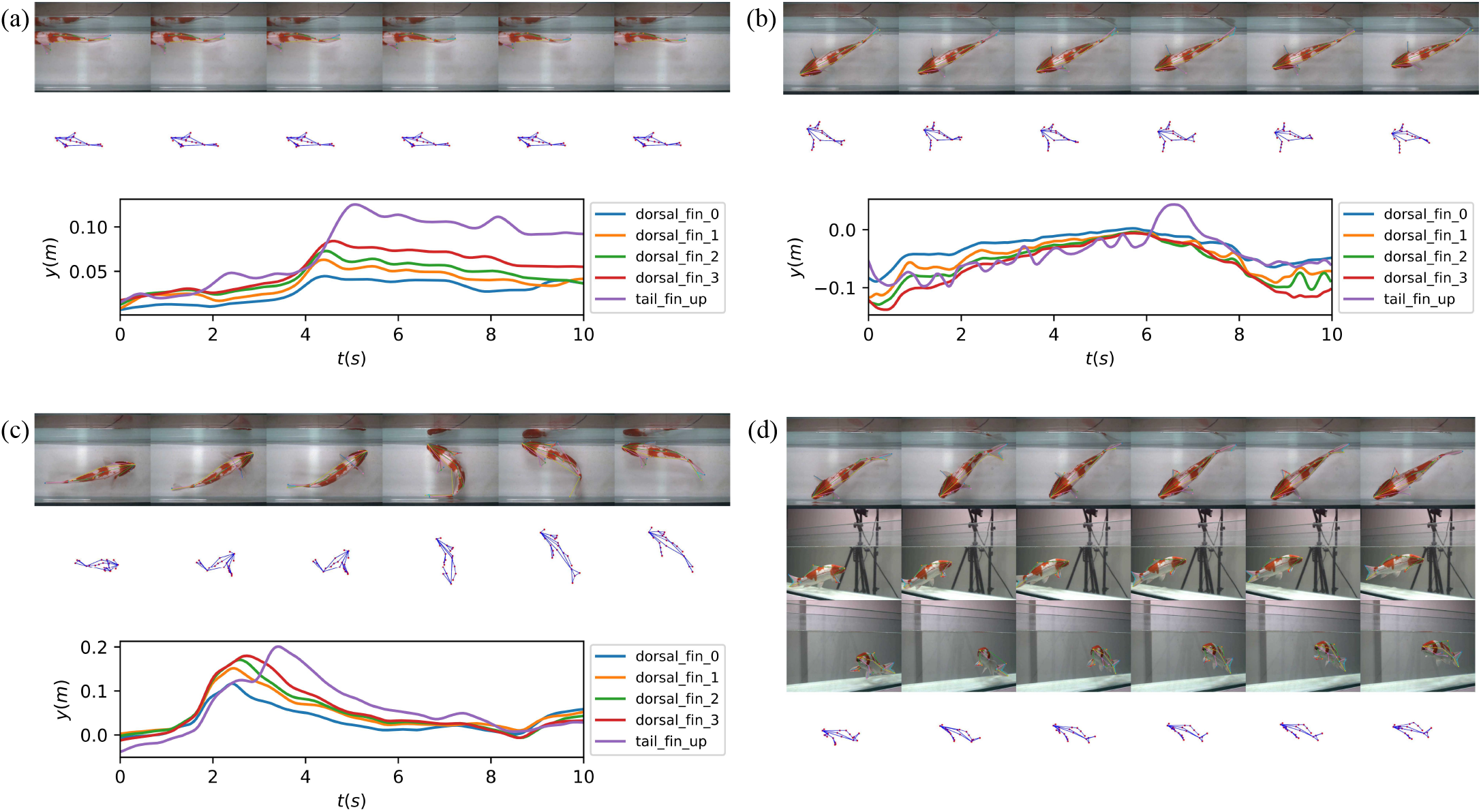
Motion feature extraction for different swimming scenarios of koi under a top-view angle, the swimming scenarios include advancing, retreating, turn, and floating up, and the motion features include 2D/3D pose estimation and displacement of five keypoints on the back of the fish in the ***y***-direction. The ***y***-direction displacement helps us understand the fish’s body wave. (a) Overlooking the koi advancing; (b) Overlooking the koi retreating; (c) Overlooking the koi advancing turning; (d) An example of 3D pose estimation from different views and a virtual perspective when the koi floats up. Different colored curves in each graph indicate the ***y***-direction displacement characteristics at different keypoints.

#### 1) Overlooking the koi advancing

The advancing state of the koi over a while is shown in Figure 7(a). The first row in the figure shows consecutive frames of the actual viewing angle, and the second row shows the 3D pose estimations from a virtual perspective. Both rows show slight bending of the fish’s body, which is consistent with the small oscillations of real fish as they swim advancing at slow speeds.

#### 2) Overlooking the koi retreating

The actual motion of the koi’s retreating state during a period and the 3D pose estimation of the virtual perspective are shown in Figure 7(b). The same keypoints of observation are used. The displacement curves of these points in the *y* direction along the time axis are shown in Figure 7(b). For retreating, the swing of the fish’s body is not significant; instead, the body has a small amplitude quasi-periodic swing. The main driving parts for fleeing are the left and right pectoral fins. Subsequent experiments will carry out a detailed analysis of this state. As a result, the fish relies much more on the left and right pectoral fins for retreating movement than advancing movement, and at a much slower rate. The cooperation between the left and right pectoral fins and the fish’s body is analyzed in detail in the following subsection.

#### 3) Overlooking the koi turning

The top view of the actual motion status of the koi turning state in a period and its 3D pose estimation are shown in Figure 7(c). The keypoints observed are the same as those in the previous two states. The displacements of the points in the *y* direction along the time axis are shown in Figure 7(c). For turning, the fish body first has a slow adjustment process, then swings wildly, and finally, after the turning is completed, the fish body swings and adjusts slightly. The primary body parts responsible for turning are the fish body and the caudal fin, while the other fins have little effect and do not provide significant driving forces. Subsequent experiments will give more detailed body-fin coordination.

#### 4) Overlooking the koi floating up

The 2D and 3D poses of the fish surfacing are shown in Figure 7(d). We could observe the 2D poses from different viewpoints and the synthesized 3D pose representations when the koi surfaced. Floating up can be viewed as moving forward in another direction, so no further analysis will be done. However, the cooperation of multiple fins plays a key role in floating up, which will be explained in detail in subsequent experiments.

### D. Body-fin co-driven for fish swimming

Most current robotic fish are caudal fin-driven, and few are multi-fin co-driven, so understanding swimming in multi-fin co-driven fish may provide solid data support for developing multi-fin co-driven robotic fish. In this section, 13 keypoints of the body, pectoral, and caudal fins were extracted using the visual perception system to analyze each fin’s synergy in the koi’s four swimming states: advancing, retreating, turning, and floating up. The process of body-fin coordination during sinking is similar to that of surfacing, and therefore, sinking is not measured.

#### 1) Body-fin co-driven for koi advancing

In Figure 8(a), we can observe the displacement curves of various keypoints of the fish body in relation to the fish head, the base of the caudal fin, and the corresponding bases of the left and right pectoral fins. These curves are presented in rows, with each row representing the *s* − *t* curves of different keypoints relative to the same body part in the *x, y*, and *z* directions. By performing Fourier analysis on these curves, we can estimate the flapping frequency.

**Fig. 8.**
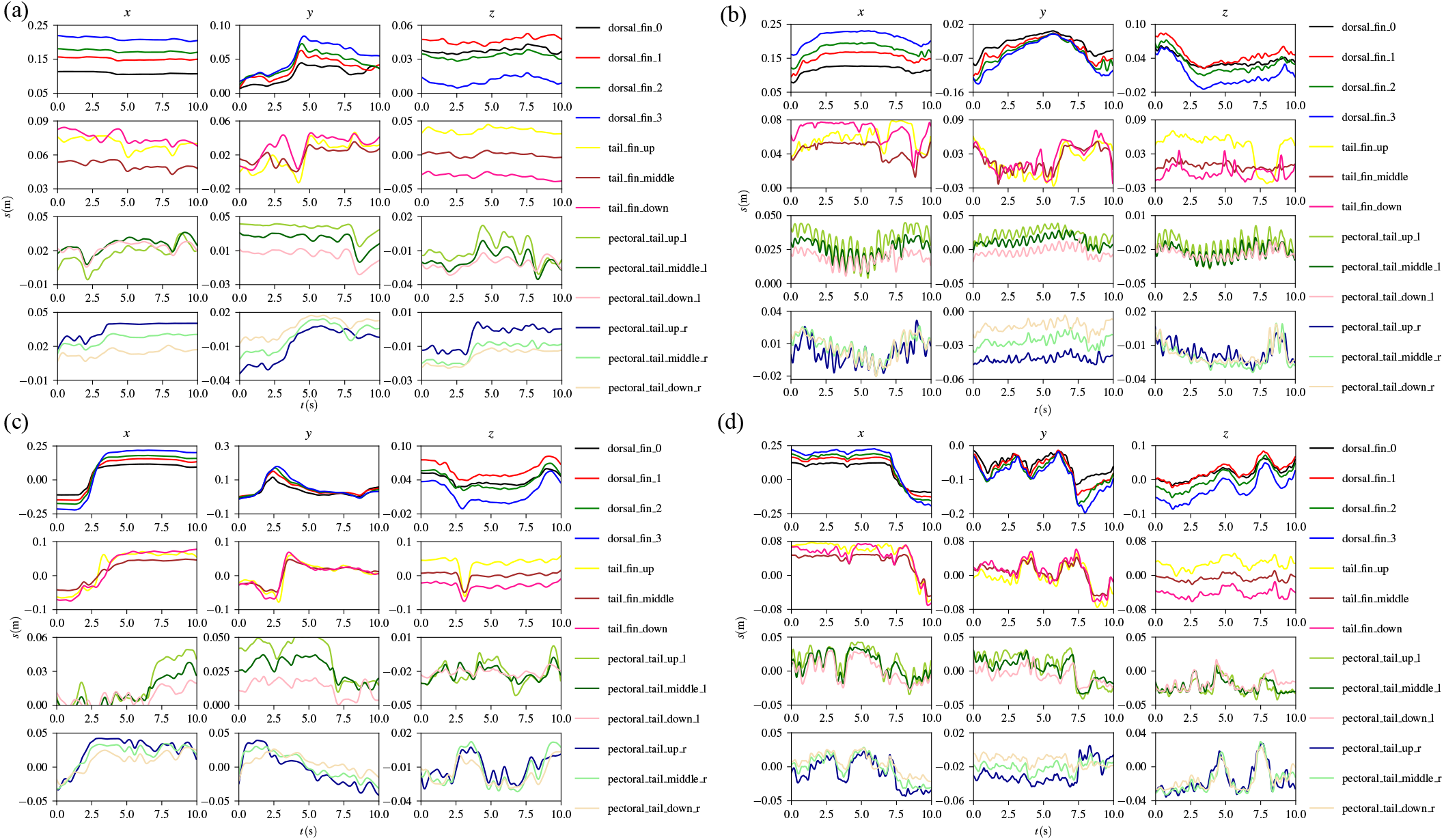
This is a characterization of the displacement of the body and fins of the fish as it swims, which is used to analyze how multiple fins work together to drive the fish to perform different swimming states. Relative displacement measurements of 13 keypoints as the koi (a) advancing; (b) retreating; (c) turning; and (d) floating up. The first row represents the change of the fish body; the second row represents the change of the caudal fin; the third row represents the change of the left pectoral fin; and the fourth row represents the change of the right pectoral fin. Different colored curves in each graph indicate the displacement characteristics at different keypoints.

Observing Figure 8(a), it becomes apparent that the koi’s body sways slowly in the *y* direction as it advances, with a maximum swing amplitude exceeding 7.5 cm. The caudal fin either sways extensively in the same direction as the fish body or slightly out of sync. Its maximum sway exceeds 3 cm, with a frequency of big swings around 0.25 Hz and small swings around 1.67 Hz. The pectoral fins flap slightly back and forth and up and down, with a maximum amplitude of over 2 cm and frequencies of 0.167 Hz and 0.89 Hz, respectively. During forward swimming, the caudal fin and body provide the main propulsion, while the pectoral fins primarily aid direction control and balance maintenance.

#### 2) Body-fin co-driven for koi retreating

From the observations in Figure 8(b), it is evident that the koi’s caudal fin does not exhibit periodic movement but shows a high deformation frequency while retreating. Meanwhile, the left pectoral fin flaps back and forth with a relatively high frequency of around 2 Hz and a maximum amplitude of 2 cm. Its up-and-down stroke amplitude is over 2 cm, and the stroke frequency is approximately 2.5 Hz. On the other hand, the right pectoral fin does not show high synchronization with the left pectoral fin, with a back-and-forth stroke frequency of about 1.32 Hz and a maximum swing amplitude of 3 cm. Its up-and-down flap amplitude is approximately 2 cm, and the flapping frequency is about 1.5 Hz. The pectoral fin oscillations are the primary driving force for the fish’s backward locomotion. In addition, the asynchronous left and right pectoral fins lead to the fish’s erratic movement as it retreats. The minimal swinging of the caudal fin and body of the fish while swimming backward also contributes to its slow swimming speed.

#### 3) Body-fin co-driven for koi turning

As depicted in Figure 8(c), the fish’s caudal fin undergoes a dramatic and high-frequency bend with the body during a turn. This entire maneuver takes approximately 4 seconds to complete, with the caudal fin swinging through a range of about 12 cm. The right pectoral fin will retract into the abdomen during a left turn. In contrast, the left pectoral fin moves away from the abdomen to maintain the fish’s balance through deformation. A sudden movement of the body primarily initiates the turning process, while the fins are responsible for balancing and controlling the direction of the turn. It is worth noting that the fish attains a remarkable speed upon completing the turn.

#### 4) Body-fin co-driven for koi floating up

Figure 8(d) presents a series of frames depicting a koi floating up. The first three rows display successive frames captured from actual observation perspectives, while the last row showcases the 3D pose estimation in the virtual view.

Observing Figure 8(d), we can note that the koi’s body swings left and right with a large amplitude of approximately 8 cm and a swing frequency of around 0.4 Hz. The caudal fin moves with the fish body, with a maximum amplitude of 5 cm and a movement frequency of approximately 0.4 Hz. On the other hand, the pectoral fins have negligible swing amplitudes, with a maximum of 2 cm. The left pectoral fin swings at about 1.5 Hz, while the right pectoral fin swings at 1.2 Hz. Additionally, the upward movement of a fish is very similar to the forward movement of a fish in that both use the body and caudal fin as the primary drivers. The difference is that the pectoral fins fluctuate less with fish uplift and the speed varies more.

### E. Representing pectoral fin morphology for fish swimming

In addition, when fish swim, each fin will deform, and the pose angle can be used to measure the bending degree of each fin. The visual result of the deformation measurement of each fin is shown in Figure 9.

**Fig. 9.**
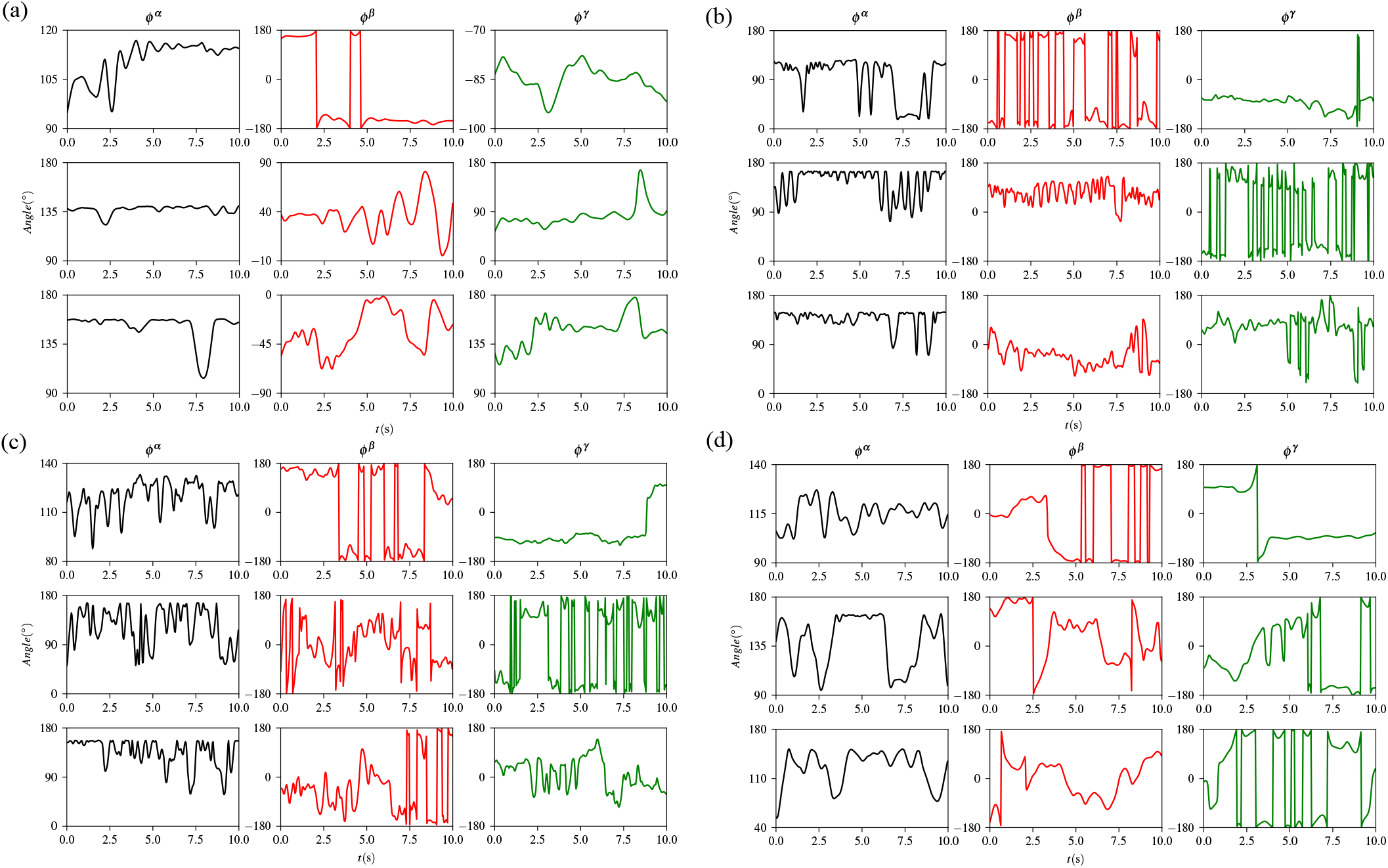
This is an angular characterization of the posture of the body and fins of a fish as it swims, and this is used to analyze how multiple fins work together to drive the fish to perform different swimming states. The deformation measurements of each fin when the koi (a) advancing; (b) retreating; (c) turning; and (d) floating up. The first row is the deformation of the caudal fin, and the second and third rows are the deformation of the left and right pectoral fins, respectively. The black curve shows the change in flexion and extension angle, the red curve shows the change in rotation angle, and the green color shows the change in torsion angle.

#### 1) Representing pectoral fin morphology for koi advancing

By observing Figure 9(a), it is evident that the koi fins undergo deformation during forward swimming. The figure’s first column displays the flexion and extension angle, with the caudal fin exhibiting a variation range of 95.0*°*∼ 116.8*°*. The left pectoral fin shows a variation range of 123.0*°* ∼ 140.4*°*, while the right pectoral fin displays a range of 103.9*°* ∼ 158.1*°*. The second and third columns of the figure showcase complex rotations, which are directional and intricate deformations that help maintain the fish’s balance and control its movement’s direction.

#### 2) Representing pectoral fin morphology for koi retreating

By observing Figure 9(b), it is evident that the koi fins undergo frequent and significant deformation during retreat. The first column of the figure displays the range of flexion and extension angles of the caudal fin (16.8*°* ∼ 126.9*°*), left pectoral fin (73.2*°* ∼ 165.*°*), and right pectoral fin (70.5*°* ∼ 149.3*°*). The figure’s second and third columns depict complex rotations involving intricate directional movements, which are crucial for maintaining the fish’s balance and controlling its retreat direction.

#### 3) Representing pectoral fin morphology for koi turning

In the turning process, the deformations of the koi fins are depicted in Figure 9(c). The first column of the figure displays the variation range of the caudal fin flexion and extension angle, which is 102.2*°* ∼ 127.1*°*, the left pectoral fin flexion and extension angle, which is 94.6*°* ∼ 164.6*°*, and the right pectoral fin flexion and extension angle, which is 53.7*°* ∼ 152.0*°*. The figure’s second and third columns show directional complex rotations that maintain the fish’s balance while controlling its turning direction.

#### 4) Representing pectoral fin morphology for koi floating up

The deformation of the koi fins during the floating-up process is shown in Figure 9(d). In the first column, the caudal fin flexion and extension angle vary between 87.9*°* ∼ 133.1*°*, while the left and right pectoral fins vary between 51.6*°* ∼ 167.0*°* and 58.1*°* ∼ 157.4*°*, respectively. These variations are much more severe than those observed in the previous motion states. The second and third columns of the figure show directional and complex rotations, which help maintain the fish’s balance and control its direction of floating up.

In summary, the complex deformation of each fin plays a crucial role in controlling the fish’s direction and position while swimming and maintaining its overall balance. Additionally, it demonstrates remarkable driving ability during the fish’s retreat.

### F. Discussion

In this section, we extracted the characteristics of koi using an visual perception system and combined these characteristics to analyze their four swimming states in still water. We analyzed koi swimming from different perspectives, with the highlight being the study of body-fin synergy in koi swimming. For koi swimming in static water with a body length of *L*, their body-fin co-driven in the four swimming states (advancing, retreating, turning, and floating up) are summarized in the following four points:

1. As the koi advances, its body sways from left to right with an amplitude of *A*_1_ less than 0.34 times its body length. The frequency of this sway falls within the range of 0.28 Hz to 0.67 Hz. Simultaneously, the tail fin also sways with the body, with an amplitude of approximately *A*_2_ and a frequency of around *f*_2_, similar to *f*_1_. Alternatively, the tail fin may sway independently with an amplitude of *A*_3_ less than 0.17 times its body length and a frequency of 3.0 ∼ 7.0*f*_1_. The pectoral fins move back and forth, or flap up and down, with frequencies of *f*_4_ = 0.5 ∼ 1.0*f*_1_ and *f*_5_ = 0.5 ∼ 1.0*f*_3_, respectively.
2. When retreating, koi rely mainly on the left and right pectoral fins for back-and-forth and up-and-down strokes, while the body and caudal fins are less affected. The left pectoral fin swings back and forth at a frequency between 2 Hz and 3 Hz, denoted by *f*_6_, while the right pectoral fin is not synchronized with the left pectoral fin at a frequency between 2 Hz and 3 Hz, denoted by *f*_7_, where *f*_6_ is not equal to *f*_7_. The asynchronous the pectoral fins resulted in the koi’s sinuous retreat route.
3. When a koi turns left or right, it achieves this through rapid and significant body and tail fin movements. The bending amplitude of the body reaches an upper limit of 0.6*L*, while the caudal fin also turns with a bending amplitude upper limit of 0.4*L*. Additionally, the pectoral fins play a role, with the right (left) fin retracting towards the fish’s belly while the left (right) fin stays away from the body.
4. When the koi floats up, the fish body swings left and right with the amplitude *A*_4_ *<* 0.27*L* and the swing frequency *f*_8_ *<*0.5 Hz. At the same time, the caudal fin swings left and right with the amplitude *A*_5_ *<* 0.17*L* and the swing frequency *f*_9_ *<*0.5 Hz, where *f*_8_ ≈ *f*_9_. The left and right pectoral fins flap up and down, while the flapping frequency of the left pectoral fin is *f*_10_ *<*2 Hz and the flapping frequency of the right pectoral fin is *f*_11_ *<*2 Hz, where *f*_10_ ≠ *f*_11_.

During all these processes, each fin undergoes complex deformation, and the deformation includes complex fin surface fluctuations. These complex deformations are mainly to maintain the balance of the fish body and control the moving directions.

To sum up, our designed visual perception system can comprehensively extract the motion characteristics of swimming fish, which can be utilized to analyze multi-fin co-driven during swimming. The exploration of multi-fin co-driven is crucial for developing multi-fin co-driven robotic fish. Additionally, the accuracy of our system’s fish swimming characterization stems from three significant advantages it has over previous vision systems:

1. We utilized multiple high-frame-rate cameras to capture fish swimming in action, compensating for the limitations of other fish-observation systems that may miss particular perspectives of raw motion and enabling us to capture even the fastest forms of motion.
2. To accurately represent the complex motions of fish swimming, we utilized an advanced animal pose estimation method. This approach addresses the shortcomings of other fish observation systems that rely on a single feature to characterize fish swimming, which often needs to be revised to capture the full range of movements.
3. We developed high-performance software that streamlines the experimental process and allows researchers to quickly obtain quantitative and qualitative representations of fish swimming quickly quantitative and qualitative fish swimming representations of fish swimming. This tool facilitates efficient and effective analysis of fish behavior.

The experimental results have effectively showcased the superior performance of the visual perception system we have designed, thus opening up avenues for further research on fish swimming. However, it is pertinent to note that despite the promising advantages of our designed visual perception system, certain areas still require further expansion and exploration to optimize its functionality and enhance its overall efficacy.

1. We are currently developing more advanced animal pose estimation algorithms. Specifically, we are focused on single-view animal 3D pose estimation and dense animal representation [36]. These advancements will help reduce equipment costs and provide richer information for studying fish surface bending.
2. It is important to note that although there have been advancements in 3D animal pose estimation, current state-of-the-art algorithms still heavily rely on 2D animal pose estimation. To address this issue, designing a novel end-to-end 3D animal pose estimation network could revolutionize the field by making pose estimation more convenient and accurate.
3. We also are exploring combining particle image velocimetry systems to create a novel representation of fish swimming with flow field and pose. This approach provides a unique perspective on fish behavior and could have promising implications for future research.

## IV. Conclusion

In this paper, a pose estimation-based experimental platform is constructed to study the movement of fish. The experimental platform can quickly capture videos of fish swimming and extract the keypoints of the fish by pose estimation algorithm and compute the motion parameters of interest. We designed the visualization software (AdamPosePlug) to enable all these steps. The experimental platform constructed in this study offers great potential for fish swimming studies.

In the future, more advanced 3D animal pose estimation algorithms can revolutionize the technical route to fish swimming analysis. In addition, this experimental platform, if combined with particle image velocimetry equipment, may be able to present fish swimming in more detail by jointly analyzing their posture and flow field data, which reminds us that animal behavior analysis techniques based on multimodal perception may be one of the essential routes for fish swimming research.

## V. Ethics STATEMENT

The experimental procedures were carried out following animal ethics approval granted by Northeast Normal University. All experimental procedures in this study were approved by the National Animal Research Authority of Northeast Normal University, China (approval number NENU-20080416) and the Forestry Bureau of Jilin Province of China (approval number [2006]178).

---

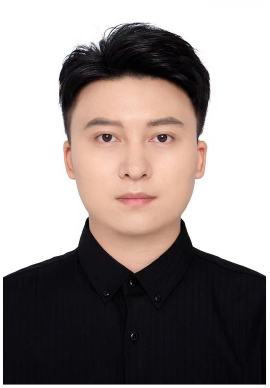

**Xin Wu** received his B.S. and M.S degrees in Electrical engineering in 2019 and in Control Science and Engineering in 2022 respectively from Northeast Normal University.

He is currently a Ph.D. candidate in School of Physics, Northeast Normal University. His research interests include computer vision, intelligent perception, biomedical measurement, and digital image procession.

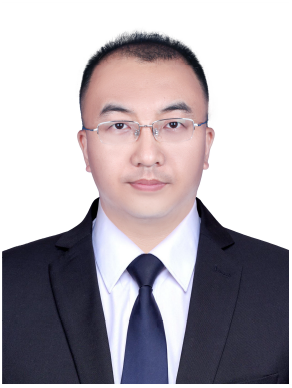

**Jipeng Huang** received the Ph.D. degree from the Changchun Institute of Optics, Fine Mechanics and Physics, Chinese Academy of Sciences, Changchun, China, in 2012.

He is currently a Professor with the School of Physics, Northeast Normal University. His research interests include the development of photoelectric detection instrument, biophysics, and digital image procession.

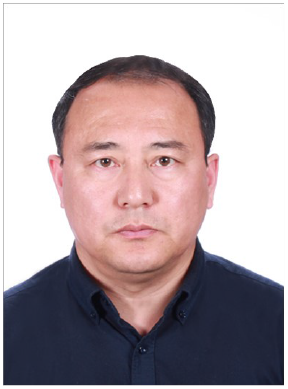

**Lianming Wang** was born in Baishan City, Jilin Province, China in 1972. He received his B.S. and M.S degrees in Electric Information Science and Technology in 1993 and in 1996 respectively from Northeast Normal University, Changchun, China and the Ph.D. degree in Mechanical and Electrical Engineering from Changchun Institute of Optics, Fine Mechanics and Physics, Chinese Academy of Sciences, Changchun, China in 2002.

He is currently a professor in the School of Marine Information Engineering, Hainan Tropical Ocean University, Sanya, China. His research interest includes computer vision, image processing and bionic underwater robot.

